# Cellular protection from H_2_O_2_ toxicity by Fv-Hsp70. Protection via catalase and gamma-glutamyl cysteine synthase

**DOI:** 10.1101/2023.02.22.529417

**Authors:** Chris Hino, Grace Chan, Gwen Jordaan, Sophia S Chang, Jacquelyn T Saunders, Mohammad T Bashir, James E Hansen, Joseph Gera, Richard H Weisbart, Robert N Nishimura

## Abstract

Heat shock proteins (HSPs), especially Hsp70 (HSPA1), have been associated with cellular protection from various cellular stresses including heat, hypoxia-ischemia, neurodegeneration, toxins, and trauma. Endogenous HSPs are often synthesized in direct response to these stresses but in many situations are inadequate in protecting cells. The present study addresses the transduction of Hsp70 into cells providing protection from acute oxidative stress by H_2_O_2_. The recombinant Fv-Hsp70 protein and two mutant Fv-Hsp70 proteins minus the ATPase domain, and minus the ATPase and terminal lid domains were tested at 0.5 and 1.0 uM concentrations after two different concentrations of H_2_O_2_ treatment. All three recombinant proteins protected SH-SY5Y cells from acute H_2_O_2_ toxicity. This data indicated that the protein binding domain was responsible for cellular protection. In addition, experiments pretreating cells with inhibitors of antioxidant proteins catalase and gamma-glutamylcysteine synthase (GGCS) before H_2_O_2_ resulted in cell death despite treatment with Fv-Hsp70, implying that both enzymes were protected from acute oxidative stress after treatment with Fv-Hsp70. This study demonstrates that Fv-Hsp70 is protective in our experiments primarily by the protein-binding domain. The Hsp70 terminal lid domain was also not necessary for protection. Cellular protection was protective via the antioxidant proteins catalase and GGCS.

## Introduction

Heat shock proteins (HSPs) are of intense interest because of their ability to protect cells from injury or death.. The proteins are divided into groups according to relative molecular mass. Of the Hsp70 group, the highly inducible Hsp70 (HSPA1), has been most studied because it is induced after stress and protect cells or organisms from heat injury. Subsequently, it has been found to protect cells from injury due to many conditions including cardiovascular related hypoxia-ischemia and oxidative stress, neurodegenerative disease, and trauma (Kampinga 2016; Kim 2020; Giffard 2008; Morimoto 2013; Beretta 2022; Dukay 2019). Overexpression of Hsp70 has been shown to protect brain from ischemic-injury in rodents (Giffard 2008). Cell survival from various stresses induce a stress response transcription factor Hsf 1 which results in the transcription and translation of many proteins and the highly inducible Hsp70 (Himanen 2022). The most studied effect of HSP induction is the inhibition of apoptosis by Hsp70 and Hsp27 (Kennedy 2014; Takayama 2003; Giffard 2008; Mehlen 1996; Srinivasan 2018). The effect of Hsp70 and Hsp27i interact with and block several proteins involved with apoptosis pathways. Hsp70 protected neonatal cardiomyocytes from ischemia/reperfusion through inhibition of p38 MAPK (Song 2020). Intracellular Hsp70 induced by prostaglandin A1 was associated with protection from rotenone-induced injury in SH-SY5Y cells. The roles of HSPs including Hsp27, Hsp70 and Hsp90 (HSPC1) were reviewed in neurodegeneration (Kampinga 2016; Beretta 2022). That review points out that the classical chaperone function of the these HSPs is suppressing toxic protein aggregation and maintaining proteostasis by facilitating abnormal protein degradation through autophagy or ubiquitin-proteosome system. Hsp27 (HSPB1) has been protective in cell culture and animal models from ischemic-injury and oxidative stress. Increased expression of Hsp27 was associated with improved cardiac function after ischemia (Ghayour-Mobarhan 2011; Behdarvandy 2020). Other protein-protein interactions between HSPs and cytoplasmic/nucleic proteins likely contribute to cellular protection.

Hypoxic-ischemic neuronal injury has been reported to be protected by mild hypothermia in animal models and human disease (Gonzalez-Ibarra 2011). Hypothermia was shown to be neuroprotective in neonatal hypoxic injury (Azzopardi 2014). An animal study showed that Hsp70 combined with valproic acid (HDAC inhibitor) and therapeutic hypothermia resulted in neuroprotection in a rat asphyxial cardiac arrest model (Oh et al 2021). In human adults there is controversy with both positive and negative outcomes with moderate hypothermia. Two Swedish studies showed no benefit from hypothermia on neurologic outcomes after cardiac arrest (Nielsen 2013; Dankiewicz 2021). Another study showed hypothermia after cardiac arrest resulted in favorable neurologic outcomes 90 days after arrest (Lasscarrou 2019). Despite the controversy, the American Academy of Neurology after review of relevant human studies recommended therapeutic hypothermia for comatose patients after cardiopulmonary arrest (Geocadin 2017; Fugate 2013). They recommended therapeutic cooling to 32-34 C for 24 hours. An *in vitro* study of H_2_O_2_ toxicity in cardiomyocytes showed protection with severe hypothermia (Diestel A et al 2011). The combination of intracellular upregulation of Hsp70 with hypothermia could theoretically result in additive prevention of cell death.

Therapeutic intracellular protein therapy is a promising therapy but has a major obstacle of crossing the cell membrane. This has been successfully circumvented by several methods called as a group, cell penetrating peptides (CPP). These include HIV-1 tat peptide, HSV VP-22, and polyarginine peptides (Lai 2005; Phelan 1998; Wadia 2005; Matsui 2003). One of the goals of our research has been to test another CPP to introduce Hsp70 intracellularly to prevent neuronal death. Although this has been accomplished by gene therapy and small molecule inducers of Hsp70, we have accomplished this by using antibody protein transduction of Hsp70 to target tissue. To accomplish this, we have exploited an antibody which passes through cell membranes. The antibody, 3E10, a mouse antinuclear antibody, was discovered by Dr. Richard Weisbart, in his study of lupus erythematosus antibodies in mouse that entered the cell nucleus without causing injury to the cell. Early studies in our laboratory collaboration resulted in the conjugation of 3E10 antibody to catalase. This construct protected primary rat neurons from H_2_O_2_ induced cell death by 73% versus 20% survival after H_2_O_2_ treatment alone (Weisbart 2000). Further studies of the antibody revealed a 30 KDa derivative of 3E10 which was more effective in passing through cell membranes (Weisbart 2004). This derivative, noted as sc3E10 or Fv, was used as a transporter through cell membranes. Transported proteins included recombinant Fv-Hsp70, which included the complete Hsp70 protein sequence. This recombinant protein, Fv-Hsp70, was used in an initial study in primary rat neurons in culture exposed to H_2_O_2_ (Hansen 2006). This resulted in 83% protection of the neurons 24 hours after exposure to H_2_O_2_ versus 41% for PBS treated controls and 46% for exogenous Hsp70. Subsequent animal studies used Fv-Hsp70 treatment after two hours of ischemic brain injury in rats (Zhan 2010). This resulted in 68% decreased infarct volume from untreated controls. This was followed by a cardiac arrest model in rabbits which shows that Fv-Hsp70 given after 40 minutes of cardiac ischemic injury resulted in marked improvement from control untreated animals (Tanimoto 2017). Both of these studies raised the question about the mechanism of Fv-Hsp70 protection of cells and whether this was due to modulating acute oxidative injury in models of ischemia followed by reperfusion. Blocking apoptosis and other cell death pathways by endogenous HSPs likely plays a role. Were other mechanisms involved? Another question was if any new construct or modification of Hsp70 within Fv-Hsp70 could be as effective as the full length Hsp70 in acute oxidative injury. This question was partially answered by previous work of engineered Hsp70 gene transfection with deletion of the ATPase site (Li 1992; Giffard 2008; Sun 2006). All of those papers demonstrated that overexpressed Hsp70 absent the ATPase site was still protective from heat shock or ischemic injury. We wanted to know whether the Fv-Hsp70 construct minus the ATPase domain continued to protect cells from injury.

This study demonstrates that two transfected Fv-Hsp70 truncated constructs were as effective in protecting cells from severe acute oxidative stress as the original Fv-Hsp70. These recombinant proteins included Fv-Hsp70 minus the ATPase domain or minus the ATPase domain and the terminal lid domain, **Figure 1A**. The observation that Fv-Hsp70 was effective in protecting cells within 15 minutes after H_2_O_2_ treatment raised the question of whether the treated cells were incapable of producing an acute protective stress response involving endogenous HSPs, specifically through heat shock transcription factor 1 (Hsf 1) induction of Hsp70 and Nrf 2 induction of heme-oxygenase 1 (HO-1). HO-1 induction has also been associated with cell survival from oxidative stress. Nrf 2 is well known to regulate the antioxidant glutathione (GSH) system and antioxidant thioredoxine system (Benarroch 2017). Inhibitors of catalase and glutathione (GSH) showed that Fv-Hsp70 significantly helped protect cells from H_2_O_2_ toxicity by protecting the activity of these antioxidative pathways without increased synthesis of Hsp70, Hsp27, or HO-1. For these studies we also utilized therapeutic hypothermia after H_2_O_2_ throughout our experiments.

**Figure 1A.**
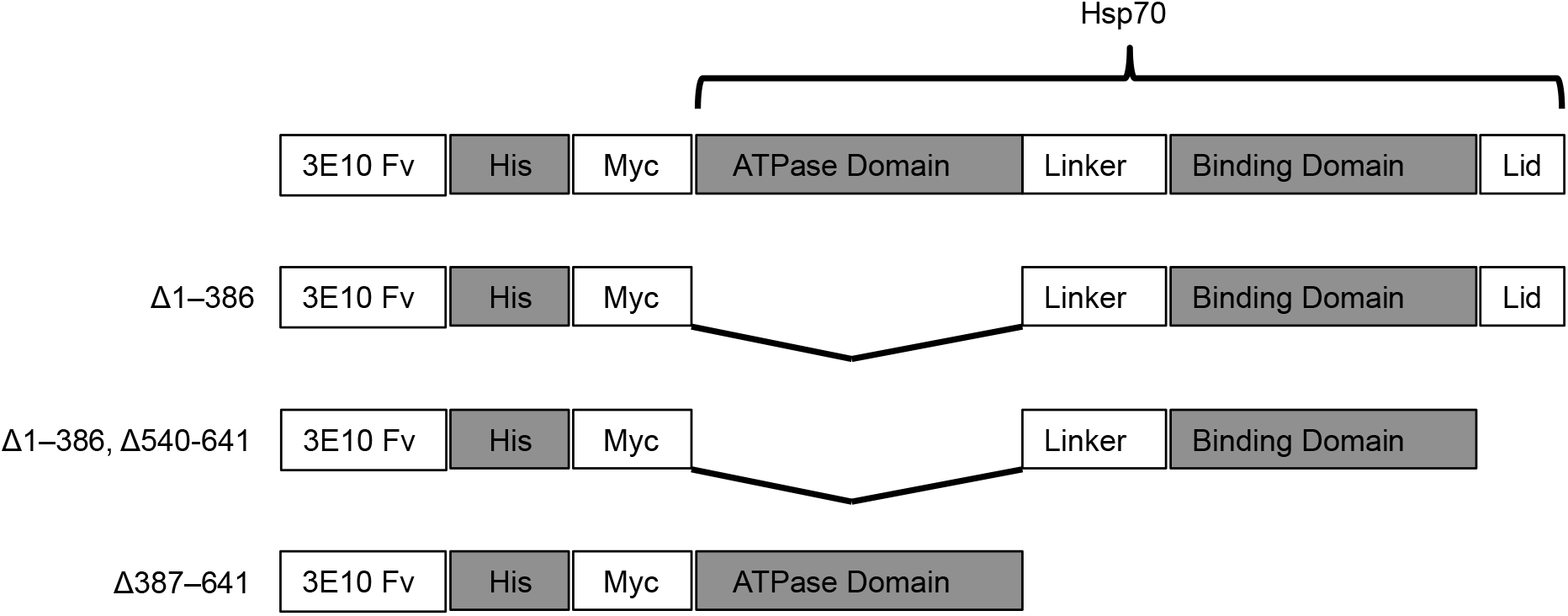
Schematic of Fv-Hs70 recombinant constructs. Fv-Hsp70, Fv-Hsp70 minus the ATPase domain, and Fv-Hsp70 minus the ATPase and Lid domains. The left margin shows the amino acids deletions resulting in the Fv-Hsp70 BD and Fv-Hsp70 BD-L.

## Materials and Methods

### Chemicals

Chemicals: 3 amino-1,2,4-triazole (3AT) (Sigma A8056), L-buthionine (S,R)-sulfoximine (BSO) (Sigma B2515). 3AT, an inhibitor of catalase, was solubilized in sterile water then added at 1 mM final concentration for 4 hours to cells prior to H_2_O_2_ treatment (Ruiz-Ojeda 2016; Koepke 2008; Smith 2007). BSO, an inhibitor of gamma-glutamylcysteine synthase (necessary for the synthesis of glutathione), was solubilized in sterile water then added at 50 uM final concentration for 4 hours to cells prior to treatment with H_2_O_2_ (Griffith 1979; Marengo 2011; Shrieve 1986). All cell culture media, PBS and chemicals were from Fisher Scientific or Sigma Chemical Company.

### Constructs of Fv-Hsp70

The original Fv-Hsp70 construct from our previous publication was used as the template for the constructs made in this study (Hansen 2006). Details of the method of producing the Fv-Hsp70 recombinant protein were published in that publication. Hsp70 protein consists of three major components. An N-terminal ATPase domain, followed by the protein binding domain (PBD) and finally the terminal lid (L) containing the terminal EEVD sequence (Mosser 2000). The EEVD sequence was described previously as necessary for protein interaction with client proteins (Mosser 2000). As demonstrated in **Figure 1A**, three recombinant proteins were produced. The original Fv-Hsp70 construct, and two deletion mutants Fv-Hsp70 absent the ATPase domain, and Fv-Hsp70 absent the ATPase and terminal lid domains. **Figure 1B** is a western immunoblot demonstrating the three constructs compared with native Hsp70 protein probed with polyclonal Hsp70 antibody.

**Figure 1B.**
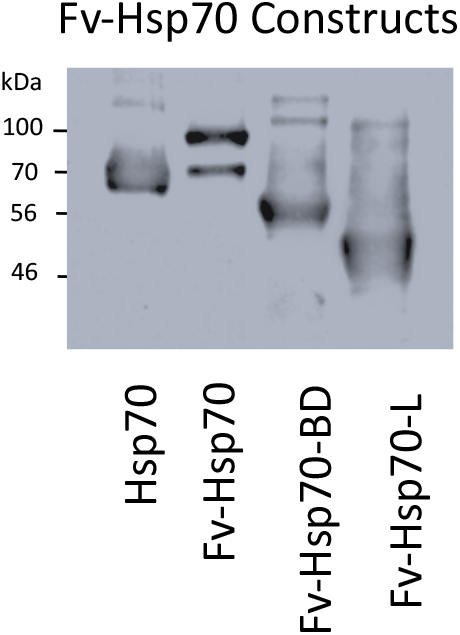
Western blot of all recombinant protein constructs in Figure 1A compared with Hsp70. Recombinant proteins 100 kD Fv-Hsp70, 56 kD Fv-70 BD, 46 kD Fv-Hsp70 BD-L are compared with Hsp70.

### Cell culture

All experiments utilizing the recombinant constructs were tested in the human neuroblastoma cell line SH-SY5Y from ATCC (CRL-2286). All experiments used undifferentiated cells. Cells were plated in 12.5 cc^2^ flasks in DMEM/F-12 medium supplemented with 10% calf serum at 37°C. When cells reached 80-90% confluence, medium was replaced with 4 ml of fresh 37°C medium. Cells were then incubated at 37°C for one hour then treated with designated final concentrations of H_2_O_2,_ then 0.5uM or 1 uM Fv-Hsp70 recombinant proteins, **Figure 2**. Control cells were treated with PBS only. The cells were then incubated overnight at 33°C (therapeutic hypothermia) and cell death was documented 24 hours after treatment. Therapeutic hypothermia was partially protective from H_2_O_2_ toxicity. Cells treated with 350 uM H_2_O_2_ at 33°C, 92.5% + 5 vs 37°C showed 98.5 % + 2, P<0.06) cell death. On the basis of that result all experiments included therapeutic hypothermia. Cells were treated with H_2_O_2_ then Fv-Hsp70 recombinant protein within 15 minutes because delaying addition of Fv-Hsp70 did not protect cells. Addition of Fv-Hsp70 after 15 minute of H_2_O_2_ treatment resulted in 90% + 4 cell death compared with 97% + 1 with H_2_O_2_ alone.

Cell death was documented by addition of 1 ug/ml propidium iodide to culture medium. Cells were then observed on Olympus IMT-2 Fluorescent Microscope with Hoffman objectives. Digital images were documented on a XCAM 8.0MP digital camera.

### Western blot

Because of the significant loss of cells secondary to H_2_O_2_ treatment in untreated controls, treated cultures were harvested 30 minutes after treatment with H_2_O_2,_ inhibitors and Fv constructs, **Figure** 4. This allowed for the acute biochemical examination of H_2_O_2_ treatment in cells before they lifted off the culture flask surface. After treatment of cells, culture medium was carefully removed by pipet and cells were washed with PBS twice and harvested in sample buffer (2% SDS, 62 uM Tris base, 5% 2 mercapto-ethanol, and 10% glycerol. Harvested cell lysates were boiled for 3 minutes. Protein content was measured using Qubit Protein Assay Kit (Thermo Scientific™). Equal aliquots of protein were loaded onto 5-20% polyacrylamide gels (Bio-Rad, Hercules, CA, USA) run for 80 minutes at 50V then blotted onto PVDF membrane (Millipore, Bedford, MA, USA) overnight at 4°C.

After transfer, blotted membranes were processed by Thermo-Fisher western blot protocol. Antibodies used for probing for specific proteins were: actin (Sigma A-2066), Hsp70 (Thermo-Fisher MA 3006), polyclonal phosphoHsp27 (Thermo-Fisher PIPA 585351), Hsp27 (Invitrogen PA5-85351), polyclonal Hsf1 (Abcam 2923), p326serHsf1 (Abcam 76076), Nrf 2 (Invitrogen PA5-38084), HO-1 (Cell Signaling 5853S), and polyclonal Nrf 2 (Cell Signaling 336495). All antibodies were used at 1:1000 dilution in blocking buffer. Developer was per the protocol for Western SuperSignal (Pierce). Developed immunoblots were imaged using LabForce Hyblot film and developed on a Konica SRX-101A X-ray film developer.

### Statistical Analysis

Determination of P values was by two-tailed Student T-test using GraphPad. Error bars represent SD.

## Results

The schematic of the Fv-hsp70 constructs is shown in **Figure 1A**. The constructs used in this study include the Fv-Hsp70 complete Hsp70 protein, Fv-Hsp70 absent the ATPase domain but includes the protein binding domain with the terminal Lid (Fv-70 BD), and the Fv-Hsp70 minus the ATPase and Lid domains (Fv70 BD-L). The Fv-Hsp70 containing only the ATPase site was not used in this study. **Figure 1B** is a western immunoblot of the Fv-Hsp70 constructs compared with 70 kD Hsp70, 100 kD Fv-Hsp70, 56 kD Fv-Hsp70 minus the ATPase domain, and 46 kD Fv-Hsp70 minus the ATPase and Lid domains.

In this study we looked at the effect of H_2_O_2_ toxicity at two final concentrations of 260 uM and 350 uM. **Figure 2A** shows the effect of 260 uM H_2_O_2_ toxicity on SH-SY5Y cells 24 hours after treatment with the Fv-Hsp70 constructs. All three constructs at two different concentrations of 0.5 uM and 1 uM significantly protected the cells from 260 uM H_2_O_2_ toxicity when compared with H_2_O_2_ treatment alone. At 350 uM H_2_O_2_ concentration the constructs showed significant protection for all Fv-Hsp70 constructs at 0.5 uM and 1 uM concentrations, while the H_2_O_2_ treated control was almost completely killed, **Figure 2B. Figure 2C** shows the effect of 350 uM H_2_O_2_ on treated, B vs control untreated cells, A. Figures 2C, 2D and 2E shows the minimal effect of H_2_O_2_ treated with 1 uM Fv-Hsp70 constructs on cells after 24 hours. Control untreated cells were almost completely killed by the 350 uM H_2_O_2_. Further experiments involved only 350 uM concentrations of H_2_O_2_ because the severe loss of cells in untreated controls made differences more significant.

**Figure 2A.**
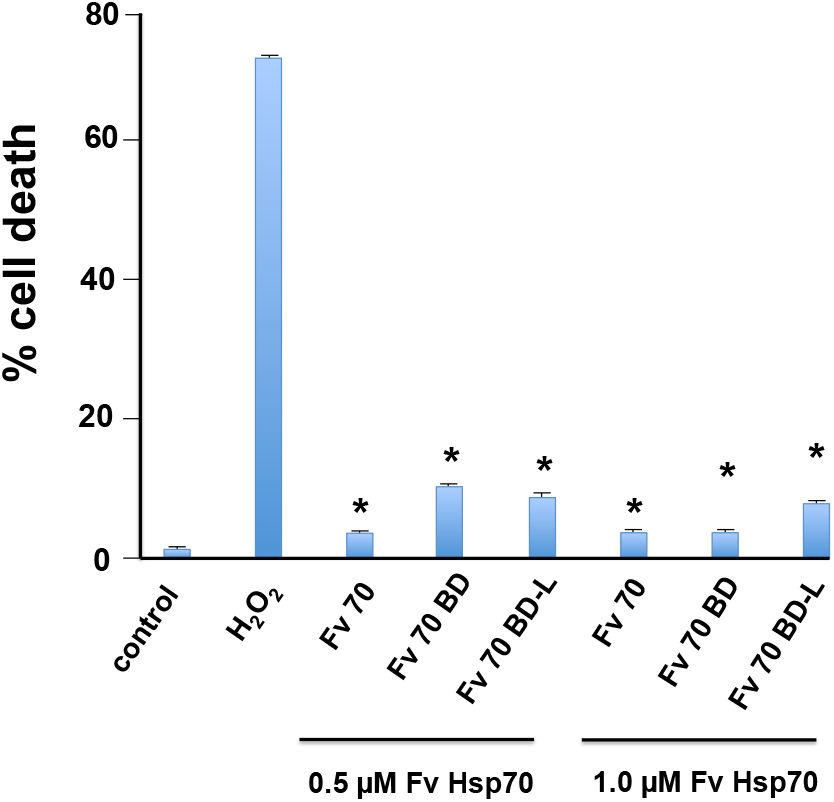
Protection of SH-SY5Y cells from 260 uM H_2_O_2_ by Fv-Hsp70 recombinant proteins. Two concentrations of Fv-Hsp70 recombinant proteins 0.5 and 1.0 uM showed the same protective effect. Protection of all Fv-Hsp70 constructs was significant (*, P < 0.01).

**Figure 2B.**
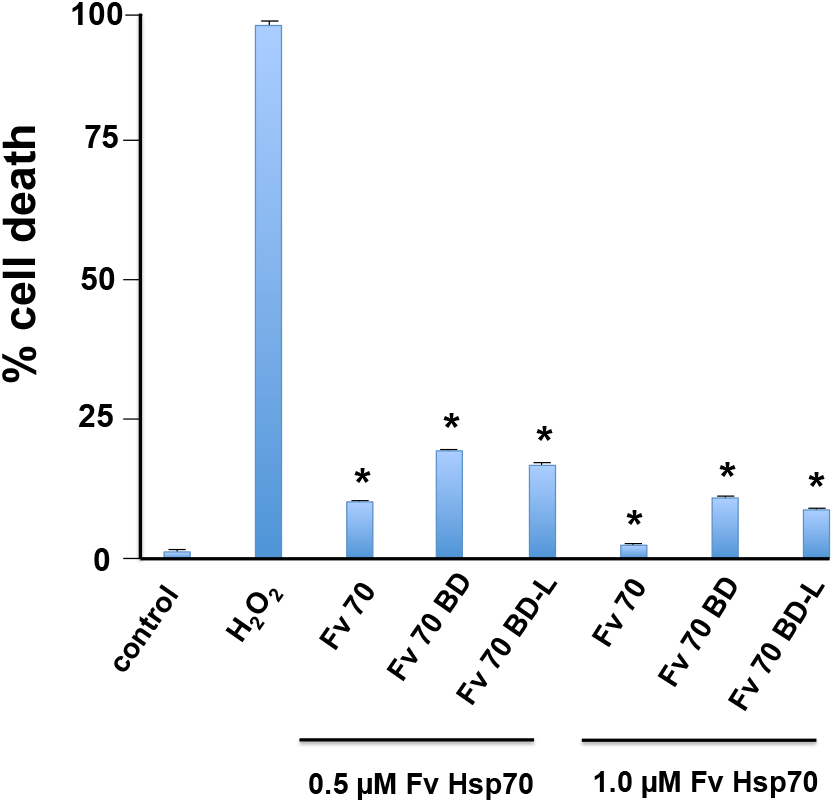
Protection of SH-SY5Y cells from 350 uM H_2_O_2_ by Fv-Hsp70 recombinant proteins. Two concentrations of Fv-Hsp70 recombinant proteins 0.5 and 1.0 uM showed the same protective effect. Protection of all Fv-Hsp70 constructs was significant (*, P < 0.01).

**Figure 2C.**
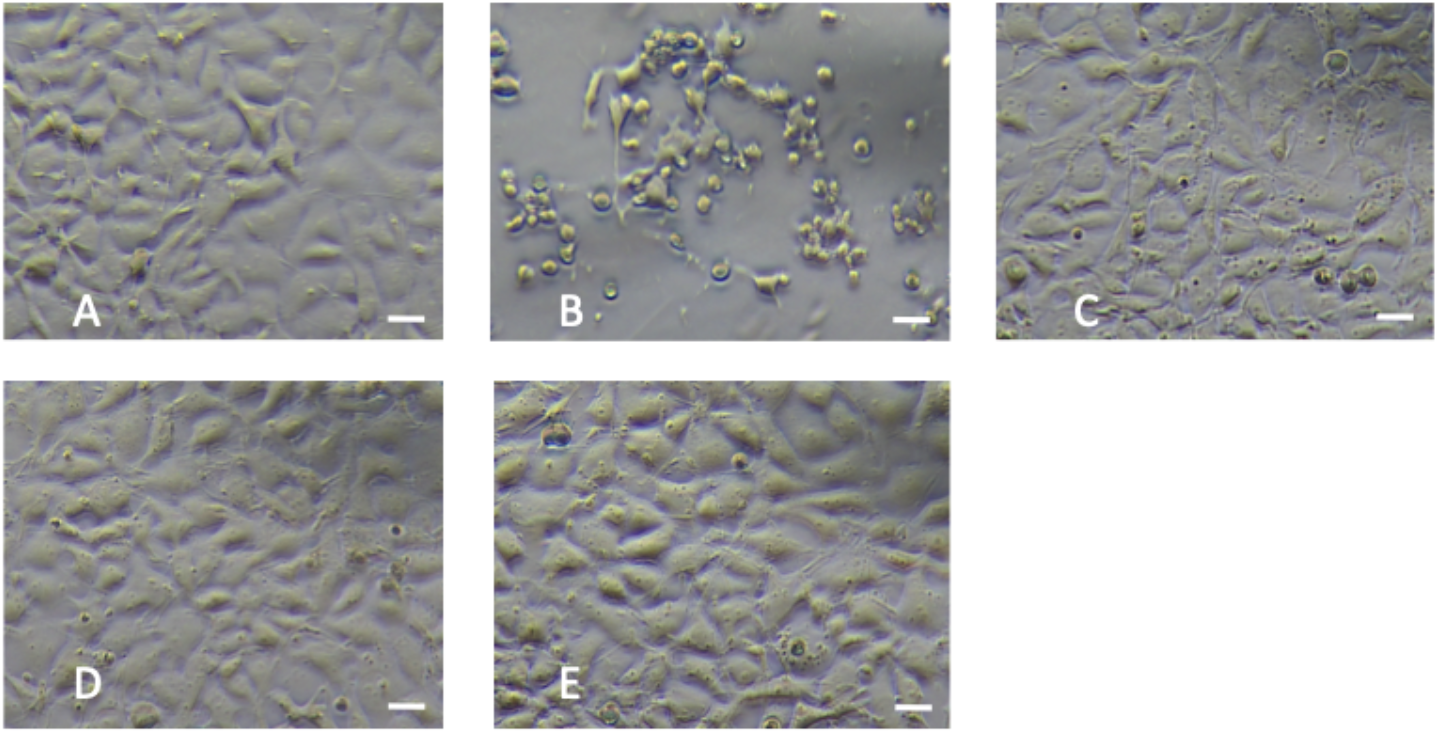
Photomicrographs of cells treated with H_2_O_2_ then Fv-Hsp70 recombinant proteins. A. Control untreated cells. B. Cells treated with 350 uM H_2_O_2_ only. C. Cells treated with 350 uM H_2_O_2_ and Fv-Hsp70. D. Cells treated with 350 uM H_2_O_2_ and Fv-Hsp70-BD+Lid. E. Cells treated with 350 uM H_2_O_2_ and Fv-Hsp70-BD. All cells were observed 24 hours after treatment. Scale bar = 20 µM.

A major question raised by our findings was the mechanism of Fv-Hsp70 cell protection from H_2_O_2_ oxidative toxicity. In previous studies when Fv-Hsp70 was delayed treating cells after H_2_O_2_ 15-30 minutes, the cells died despite treatment with the Fv-Hsp70. We reasoned that the acute protective effect of the Fv-Hsp70 recombinant protein constructs were most likely the result of interaction with nondenatured antioxidative proteins such as catalase and those involved in the production of antioxidant glutathione. If Fv-Hsp70 interaction with either antioxidant pathway was protecting cells from death then inhibiting these pathways would result in cell death. Fv-Hsp70 would be less protective. Inhibitor of catalase, 3-amino-1,2,4-triazole (3AT) and gamma-glutamylcysteine synthase (GGCS) inhibitor, L-buthionine (S,R)-sulfoximine (BSO), were used to pretreat cells prior to H_2_O_2_ and Fv-Hsp70 treatment. BSO inhibited the synthesis of the antioxidant glutathione. Cells were treated with 50 uM BSO or 1 mM 3AT for 4 hours prior to 350 uM H_2_O_2_ and 1 uM Fv-Hsp70 (complete Hsp70 protein). **Figure 3** shows the effect of BSO inhibition of GGCS resulting in significant cell death after H_2_O_2_ exposure despite Fv-Hsp70 treatment. Inhibition of catalase with 3AT also resulted in significant cells death from H_2_O_2_ and Fv-Hsp70 treatment but to a lesser extent than cells treated with BSO. Inhibition of both catalase and GGCS also resulted in almost complete loss of cells when treated with H_2_O_2_ and Fv-Hsp70. The experiments were repeated three times and the **Figure 3A and 3B** are representative of results. These data provide evidence that antioxidative enzymes may interact with Fv-Hsp70 to provide cell protection due to H_2_O_2_. **Figure 3B** shows cells treated with inhibitors of catalase and GGCS treated with H_2_O_2_ and Fv-Hsp70. Cells treated with the antioxidative enzyme inhibitor BSO resulted in more cell death than AT treated cells. The round cells noted treated with BSO or 3AT were dead cells, **Figure 3B**.

**Figure 3A.**
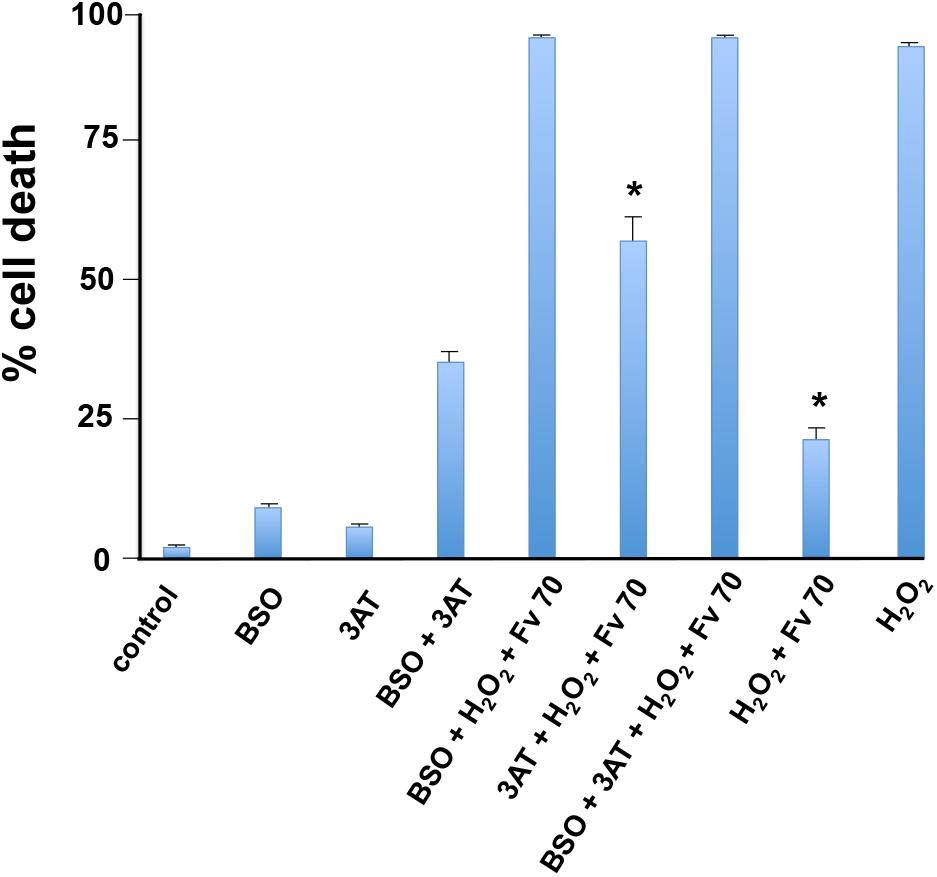
Cells treated with inhibitors of catalase (3AT) and gamma-glutamyl cysteine synthase (BSO) then H_2_O_2_ and Fv-Hsp70. Inhibitors treatment 4 hours prior to H_2_O_2_ and Fv-Hsp70 neutralized the protective effect of Fv-Hsp70. Inhibition of GGCS caused more cell death than inhibition of catalase. Treatment with Fv-Hsp70 was significantly protective from H_2_O_2_ treatment. Treatment with 3AT + H_2_O_2_ and Fv-Hsp70 was also significantly protected from H_2_O_2_ treatment only (*, P < 0.01).

**Figure 3B.**
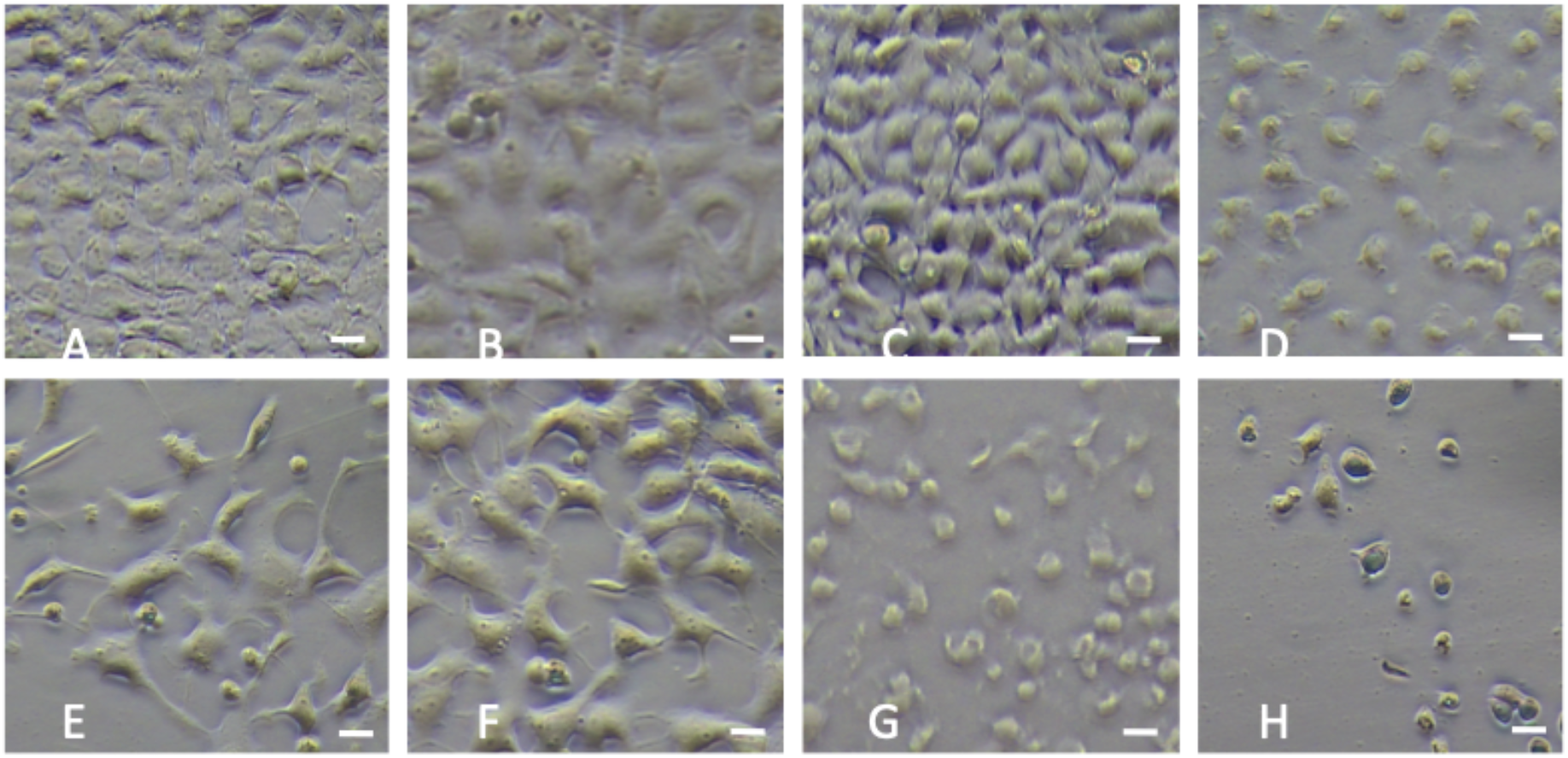
Cells treated with inhibitors 3AT and BSO then H_2_O_2_ and Fv-Hsp70. A. control untreated cells. B. BSO treated control. C. 3AT treated control. D. BSO then H_2_O_2_ and Fv-Hsp70. E. 3AT treated then H_2_O_2_ and Fv-Hsp70. F. H_2_O_2_ and Fv-Hsp70 treated. G. BSO and 3AT then H_2_O_2_ and Fv-Hsp70 treated. H. H_2_O_2_ treated only. Scale bar = 20 µM.

We then hypothesized that the heat shock protein response to H_2_O_2_ did not contribute to the cell protection from acute oxidative stress due to H_2_O_2_ because of the brief time window for efficacy of Fv-Hsp70 antiapoptotic protection. We also felt that Nrf 2 induction of HO-1 which is reported to be protective from oxidative stress was also inadequate. To test whether either HSPs or HO-1 induction was inadequate to protect cells we treated cells with inhibitors of catalase and GGCS then H_2_O_2_ and Fv-Hsp70 (complete Hsp70 protein) for 30 minutes then harvested cells and probed for HSPs and HO-1. We chose the 30 minute time point after treatment because the loss of cells after that time would make analysis for proteins difficult and we considered that within the acute stage of toxicity. **Figure 4A** shows that cells treated with BSO or 3AT did not induce a significant heat shock response as Hsp70 and Hsp27/pHsp27 did not show induction. Hsf 1 was phosphorylated indicating activation of the stress response but did not result in significant induction of Hsp70 or Hsp27/pHsp27. All cells treated with H_2_O_2_ with or without BSO or 3AT and Fv-Hsp70 showed no significant differences in Hsp70 or Hsp27 induction. However, all cells treated with H_2_O_2_ with or without inhibitors and Fv-Hsp70 show phosphorylation of Hsp27. These data suggested that newly synthesized Hsp70 or Hsp27 was not involved with the acute protection from oxidative stress. The role of phosphorylated Hsp27 was not clear. Another antioxidant pathway induced by oxidative stress is controlled by Nrf 2 induction. Activation of the Nrf 2 pathway results in synthesis of heme-oxygenase 1 (HO-1) and many antioxidant proteins reported to be protective in response to oxidative injury (Benarroch 2017; Biswas (2014). **Figure 4B**) shows that Nrf 2 was not induced by acute treatment with H_2_O_2_ and that subsequent increased HO-1 induction was not seen. The effect of inhibitors BSO and 3AT did not affect induction. There was no difference in treated controls compared with experimental groups. We concluded that Nrf 2 was also not involved in the acute protection from severe oxidative stress of H_2_O_2_.

**Figure 4A.**
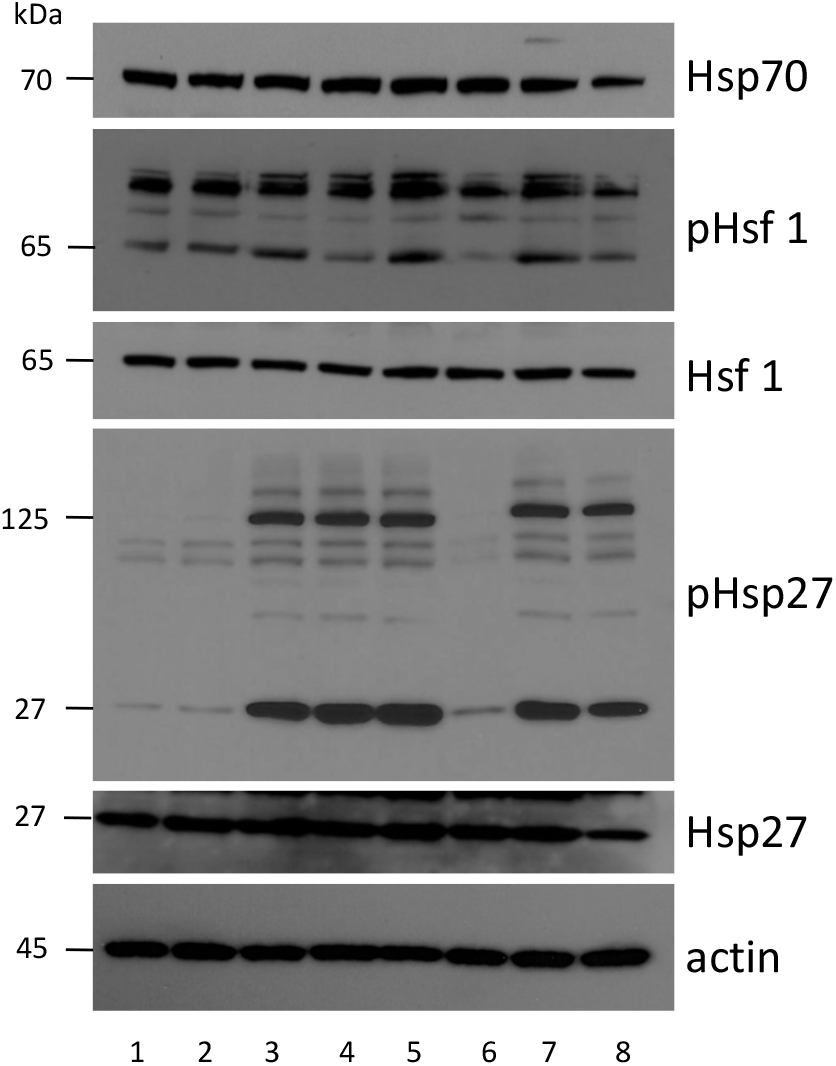
Western blot analysis of the Hsp70 and Hsp27 induction after treatment with BSO, 3AT, with or without H_2_O_2_ and Fv-Hsp70 treatment. All westerns were analyzed from a single experiment. The results are representative of three experiments. 1. Control untreated. 2. BSO treatment only. 3. H_2_O_2_ only. 4. BSO + H_2_O_2_ + Fv-Hsp70. 5. H_2_O_2_ + Fv-Hsp70. 6. 3AT only. 7. 3AT + H_2_O_2_ + Fv-Hsp70. 8. H_2_O_2_ + Fv-Hsp70.

**Figure 4B.**
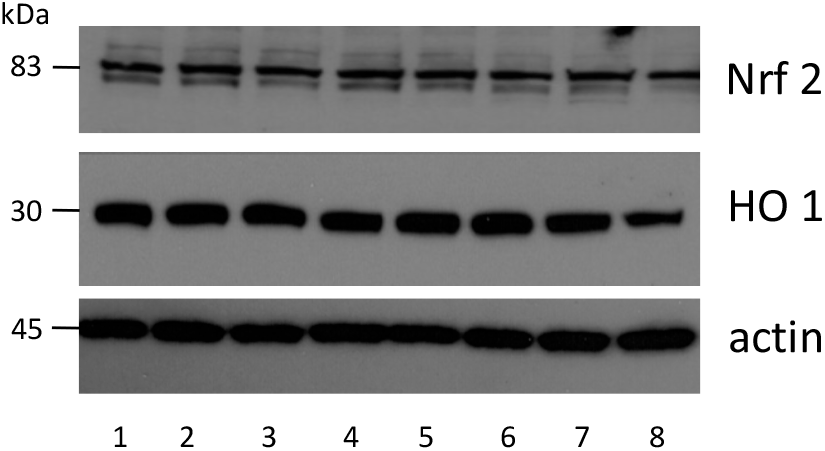
Western blot analysis of Nrf 2 and HO-1 induction after treatment as noted in Figure 4A. These were samples from the same experiment in 4A. Actin control is from Figure 4A. The results are representative of three experiments. 1. Control untreated. 2. BSO treatment only. 3. H_2_O_2_ only. 4. BSO + H_2_O_2_ + Fv-Hsp70. 5. H_2_O_2_ + Fv-Hsp70. 6. 3AT only. 7. 3AT + H_2_O_2_ + Fv-Hsp70. 8. H_2_O_2_ + Fv-Hsp70.

## Discussion

The data demonstrates that Fv-Hsp70 and derivatives were cytoprotective from oxidative stress. In addition, the acute protective effect is likely due to protecting antioxidative proteins from denaturation during H_2_O_2_ exposure. Truncated Hsp70 recombinant protein absent the ATPase domain was protective against heat stress when genetically transfected into rodent cells (Li et al 1992; Giffard 2008). Overexpression of Hsp70 mutants including an ATPase deficient point mutation and Hsp70-381-640 deletion of ATPase domain mutant protected astrocyte mitochondrial glucose deprivation stress and middle cerebral artery occlusion injury followed by 24 hours of reperfusion (Sun et al 2006; Ouyang et al 2006). A similar study of Hsp70-ATPase deleted and Hsp70-EEVD deleted mutant cells failed to protect cells from stress-induced apoptosis (Mosser et al 2000). All of those studies involved overexpression of mutated Hsp70 in cells.

The present study used an antibody, single-chain Fv recombinant protein, Fv-Hsp70, and mutants minus the ATPase domain and the terminal lid domain including the EEVD terminal peptide. These different recombinant proteins were introduced to stressed cells after acute oxidative stress by H_2_O_2_. Significant protection was provided by all three recombinant proteins including the recombinant protein minus the terminal -EEVD peptide. This was surprising as EEVD was reported to be essential for protein-protein interaction (Mosser et al 2000). The intracellular delivery of Hsp70 has also been accomplished using HIV-1 TAT transduction domain. The recombinant protein, TAT-Hsp70 has shown efficacy in several pathological conditions including hypoxia-ischemia (Doeppner et al 2009; Lai et al 2005; Doeppner et al 2013; Doeppner et al 2012), Parkinson’ s disease models (Nagel et al 2008), spinal cord ischemic injury (Kim et al 2019), lung injury (Lyons et al 2015), neuronal differentiation (Kwon et al 2019). Another TAT-Hsp70-2 protein was also produced with reported neuroprotective properties (Cappellett 2017).

A major difference between Fv-Hsp70 and TAT-Hsp70 is the route of intracellular transduction. Fv-Hsp70 enters the cell passively through the ENT 2 nucleoside salvage pathway (Hansen 2007). TAT-Hsp70 enters the cell by endocytosis. Endocytosis requires energy expenditure while Fv-Hsp70 enters passively not requiring ATP. This latter characteristic is particularly relevant in treatment of injured tissue lacking ATP production as in ischemia or trauma. Another differing characteristic is the localization within the cell. Fv-Hsp70 initially locates in the nucleus and TAT-Hsp70 to the cytoplasm. Localization of the recombinant constructs to target tissue is also important. Fv-Hsp70 localizes with DNA fragments as in injured tissue or tumors (Weisbart 2015). TAT-Hsp70 lacks a tissue targeting characteristic. In addition, there is evidence that the TAT peptide induces an inflammation response in animals which is not optimum as a transfection vehicle (Toborek 2003).

Heat shock proteins including Hsp70 and the small heat shock proteins including Hsp27 have been associated with cell survival and protection. Both Hsp70 and Hsp27 are known to be involved in survival through regulation of apoptosis (Mehlen 1996; Giffard 2008; Kennedy 2014; Takayama 2003; Srinivasan 2018). This study looked at the acute effect of protection by Fv-Hsp70 from oxidative stress by H_2_O_2_. The rationale for looking at the antioxidative proteins was due to the rapid cell death induced by H_2_O_2_ if treatment was delayed greater than 15 minutes. This time frame may be too brief for the synthesis of protective HSPs or Nrf 2 induced protective proteins. We chose inhibitors of representative antioxidative enzymes catalase and GGCS. The latter is essential for synthesis of glutathione (GSH) (Seo 2004). Glutathione related enzyme activity is modulated by Hsp70 (Guo 2007). Our findings implicate catalase and GGCS in cytoprotection by Fv-Hsp70 from acute oxidative stress. A shortcoming of using inhibitors are off-site effects which we did not investigate. Using siRNA for catalase and GGCS were not used due to the rapid growth of the cells and the unknown half-life of both enzymes. In **Figure 3A** we noted mild -moderate cell death with the inhibitors without further treatment indicating that both enzymes are likely essential in protection from oxidative stress. Further investigation of our hypothesis that induction of HSPs or Nrf 2 was insufficient for protection was supported in **Figure 4A** and **4B**. The western immunoblot showed minimal or no induction of endogenous Hsp70, Hsp27, Nrf 2, and HO-1. A possible reason for the lack of induction of these proteins is hypothermia (Lee 2017). In that study hypothermia caused decreased HSP expression after hypoxia-ischemia of brain. The protein analysis in **Figure 4** was performed 30 minutes after treatment and unlikely to show significant reduction of protein synthesis due to therapeutic hypothermia. Another unanswered question is why endogenous HSPs did not protect cells from the inhibition of enzymes and oxidative stress. Our data does not reveal how Fv-Hsp70 might interact with catalase or GGCS. Was the initial localization of Fv-Hsp70 responsible for the protective effect? The protective effect of Fv-Hsp70 construct minus the lid domain was surprising since this domain has been implicated in protein-protein interactions (Mosser 2000). Hsp70 can be oxidized, and does this affect the endogenous HSPs or make them less effective (Grumwald 2014). If this is true then why was Fv-Hsp70 still effective? This study did not investigate the specific interactions of Fv-Hsp70 with catalase and GGCS. Also, what could be the role of a second protein binding domain in Hsp70 in protection (Lee 2017). Finally, though literature supports therapeutic hypothermia after cardiac arrest, it is unlikely to be a major factor in protection from acute oxidative stress in this study as the inhibitors of antioxidant proteins demonstrated severe cell death similar to only H_2_O_2_ treated cells.

Criticisms of this study include lack of animal verification of findings and studies only involved one human cell line. This study did not investigate specific apoptotic or ferroptosis pathways of cell death. The study only looked at a 24 hour window which may be insufficient to see if other pathways of cell death are activated. Though implied by inhibitors, this study did not show that Fv-Hsp70 had a direct protective protein-protein association with catalase or GGCS. The protective effect of Fv-Hsp70 may have been through a cellular protein or pathway not studied. Fv-Hsp70 may interact with many non-canonical proteins not yet described (Johnson and Gestwicki 2022). Fv-Hsp70 may be protecting catalase and GGCS prior to H_2_O_2_ damage and not after damage as suggested by the activity of Hsp70 homolog DnaK in E.coli (Santra 2018). This might explain the delay in Fv-Hsp70 treatment resulting in cell death by H_2_O_2_. We did not test the two deletion mutants with the antioxidant inhibitors and H_2_O_2_ toxicity. It is possible that they could protect cells from oxidative stress through a different mechanism.

## Conclusions

In summary, the present study supports Fv-Hsp70 and deletion mutants of Hsp70 in protection of cells from acute oxidative stress. In addition, the acute toxic effect of H_2_O_2_ may be partially prevented by the protein-protein interaction of Fv-Hsp70 with catalase and GGCS. Future studies in animal models might support these findings and discover new pathways involved in cellular protection from acute oxidative stress.

## Abbreviations

(HSPs): heat shock proteins
(HO-1): heme-oxygenase
(GGCS): gamma glutamyl cysteine synthase
(Hsp70): HSPA1
(Hsp27): HSPB1
(3AT): 3-amino-triazole
(BSO): buthionine sulfoxime

